# Confinement Discerns Swarmers from Planktonic Bacteria

**DOI:** 10.1101/2020.08.30.274316

**Authors:** Weijie Chen, Neha Mani, Hamid Karani, Hao Li, Sridhar Mani, Jay X. Tang

## Abstract

Powered by flagella, many bacterial species exhibit collective motion on a solid surface commonly known as swarming. As a natural example of active matter, swarming is also an essential biological phenotype associated with virulence, chemotaxis, and host pathogenesis. Physical changes like cell elongation and hyper flagellation have been shown to accompany the swarming phenotype. However, less noticeable, are the contrasts of collective motion between the swarming cells and the planktonic cells of comparable cell density. Here, we show that confining bacterial movement in designed dimensions allows distinguishing bacterial swarming from collective swimming. We found that on a soft agar plate, a novel bacterial strain *Enterobacter* sp. SM3 exhibited different motion patterns in swarming and planktonic states when confined to circular microwells of a specific range of sizes. When the confinement diameter was between 40 μm and 90 μm, swarming SM3 formed a single swirl motion pattern in the microwells whereas planktonic SM3 showed multiple swirls. Similar differential behavior is observed across a range of randomly selected gram-negative bacteria. We hypothesize that the “rafting behavior” of the swarming bacteria upon dilution might account for the motion pattern difference. We verified our conjectures via numerical simulations where swarming cells are modeled with lower repulsion and more substantial alignment force. The novel technical approach enabled us to observe swarming on a non-agar tissue surface for the first time. Our work provides the basis for characterizing bacterial swarming under more sophisticated environments, such as polymicrobial swarmer detection, and *in vivo* swarming exploration.

## Introduction

Motility is an essential characteristic of bacteria. Although energy-consuming, it provides high returns, enabling cells to uptake nutrients efficiently and escape from noxious environments(Webre, Wolanin, & Stock, 2003). In a host environment, bacterial motility is an essential phenotype that intimately relates to virulence through complex regulatory networks(Josenhans & Suerbaum, 2002). Swimming and swarming are two common motility phenotypes mediated by flagella. Whereas the planktonic phenotype defines individual bacteria’s motility, a collective movement powered by rotating flagella(Kearns, 2010a) on a partially solidified surface defines swarming(Partridge & Harshey, 2013). In swarming, bacteria utilize their flagella to navigate, two-dimensionally, through a medium and acquire necessary molecules to maintain homeostasis and overall survival(N. C. Darnton, Turner, Rojevsky, & Berg, 2010). Morphological changes like cell elongation may or may not occur in all swarming bacteria(Michaels & Tisa, 2011). Thus, concentrated swimming bacteria are often called “a swarm of bacteria” without requiring precise identification of swarming motility, per se. Nevertheless, microbiologists believe that swarming and swimming are fundamentally different motility types. For instance, studies found that compared with swimming cells, the requirement for flagella torque is higher for swarming *B. subtilis*(Hall, Subramanian, Oshiro, Canzoneri, & Kearns, 2018); swarming *E. coli* remodel their chemotaxis pathway(Partridge, Nhu, Dufour, & Harshey, 2019); and in swarming *P. aeruginosa,* both the production of virulence factors and antibiotic resistance increase(Overhage, Bains, Brazas, & Hancock, 2008). A recent study has demonstrated a medically relevant distinction between swarming and swimming: a particular strain of swarming Enterobacter protect against mice intestinal inflammation while their swimming counterparts could not(Chen et al., 2020). The evidence to date that shows swarming is different from swimming comes mostly from biological data(Kearns, 2010b). However, precise biophysical visualization and quantitation of these differences are lacking. In this report, using *Enterobacter* sp. SM3, which is a novel strain that possesses both swimming and swarming motilities, we show distinct biophysical characteristics between these two types of motility under confined, circular geometry of a particular confinement size range.

Studies have shown that geometric constraints have a profound influence on patterns of microswimmers’ collective motion. For example, these constraints may create mesoscopically or macroscopically coherent structures such as swirls and jets(Theillard, Alonso-Matilla, & Saintillan, 2017; Wioland, Lushi, & Goldstein, 2016; Wioland, Woodhouse, Dunkel, & Goldstein, 2016). Circular confinement, in particular, could stabilize a suspension of motile bacteria into a spiral vortex(Beppu et al., 2017; Lushi, Wioland, & Goldstein, 2014; Nishiguchi, Aranson, Snezhko, & Sokolov, 2018; Wioland, Woodhouse, Dunkel, Kessler, & Goldstein, 2013). Here, we compare the behaviors of bacteria in swarming and planktonic states under quasi-two dimension (quasi-2D) circular confinement. Many species of bacteria show distinctive motion patterns while confined. This characteristic may lead to future diagnostic applications since there are established associations between bacterial swarming and virulence pathology(Lane, Alteri, Smith, & Mobley, 2007; Overhage et al., 2008).

## Results

### Swarming *Enterobacter* sp. SM3 forms large single swirls

A novel bacterial strain *Enterobacter* sp. SM3 (NCBI BioProject PRJNA558971), isolated in 2014 from mice with colitis induced with dextran sulfate sodium (DSS), has been previously studied for motility(Araujo, Chen, Mani, & Tang, 2019) and host phenotype(Chen et al., 2020). SM3 expands rapidly on 0.5% agar with the collective motion of multilayers of cells at the edge. We mounted a PDMS chip containing circular microwells on the agar so that bacteria in confinement could rotate for more than 3 hours (details with illustration in Methods). Under confinement in circular wells in the diameter range of 31-90 μm, swarming SM3 shows single swirls. In contrast, SM3 planktonic cells concentrated from the liquid medium form mesoscale vortices (multiple swirls) in the same size range, except for the smallest well diameter of 31 μm. A clear difference is shown at the well diameter of 74 μm (Fig. 1A-D, Movie S1 & S2). This striking difference persists in several well depths, except that the concentrated cells yield small but non-zero vortex order parameters (VOPs, defined as illustrated in Fig. 1E) in deeper wells, instead of nearly zero VOPs in shallow wells (Fig. 1F).

**Figure 1.**
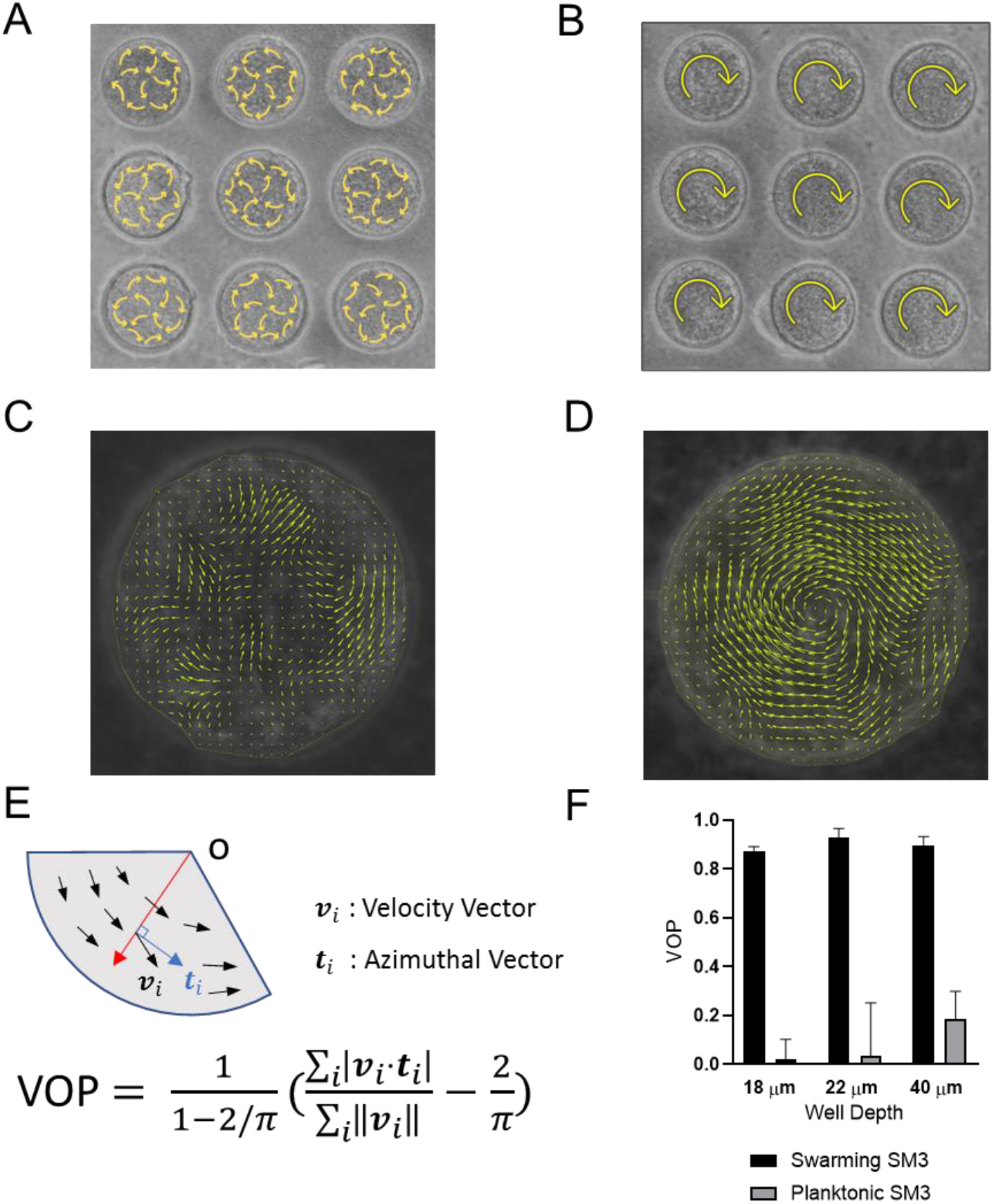
Swirls of *Enterobacter* sp. SM3 under circular confinement. (A-B) Motion pattern of concentrated planktonic (A) and swarming (B) SM3 in the PDMS microwells of 74 μm in diameter. Circular arrows indicate the direction of bacterial collective motion. (C-D) Velocity field of concentrated planktonic (C) and swarming (D) SM3 in a single microwell. (E) Illustration of how vortex order parameter (VOP) is defined. ļoļ denotes the absolute value while ļļoļļ denotes the Euclidean norm. (F) VOP of swarming and swimming SM3 in 74 μm microwells of 3 different depths. The sample size n = 5 for each group and data are represented as mean and standard deviation (±SD).

The confinement well diameter has a strong influence on the motion pattern in the wells. In smaller wells like 31 μm in diameter, even concentrated planktonic SM3 forms a single vortex (Fig. 2A), whereas in larger wells, such as ones of 112 μm in diameter, swarming SM3 also breaks into mesoscale vortices (Fig. 2B). The phase diagram shows a single swirl in small confinement for both phenotypes of SM3. The patterns diverge as confinement size increases, but they converge towards multiple swirls as confinement size reaches 144 μm and above (Fig. 2C). To further compare the dynamics of the confined swarming and swimming SM3, the spatial correlation of the velocity field was calculated for d = 90 μm (where the motion patterns differ for swarming and planktonic SM3) and for d = 500 μm (where both motilities show mesoscale vortices) (see Methods). We computed the correlation function for the inscribed square within a well, which shows the extent to which the velocity at an arbitrary location correlated with the velocity at a distance of Δr away from that location. In 90 μm wells, swarming SM3 velocity correlates positively or negatively throughout the whole well (negative values have resulted from the opposite sides of a single swirl). In contrast, the swimming velocity of planktonic cells of comparable concentration does not correlate once Δr > 25 μm (Fig. 2D). However, in a large open space where both swarming and swimming SM3 break into small vortices, the correlation functions look similar. The characteristic length as the curve first crosses Cr(Δr) = 0, which also represents the size of the mesoscale vortices of planktonic and swarming SM3 is 27 μm and 33 μm, respectively (Fig. 2E).

**Figure 2.**
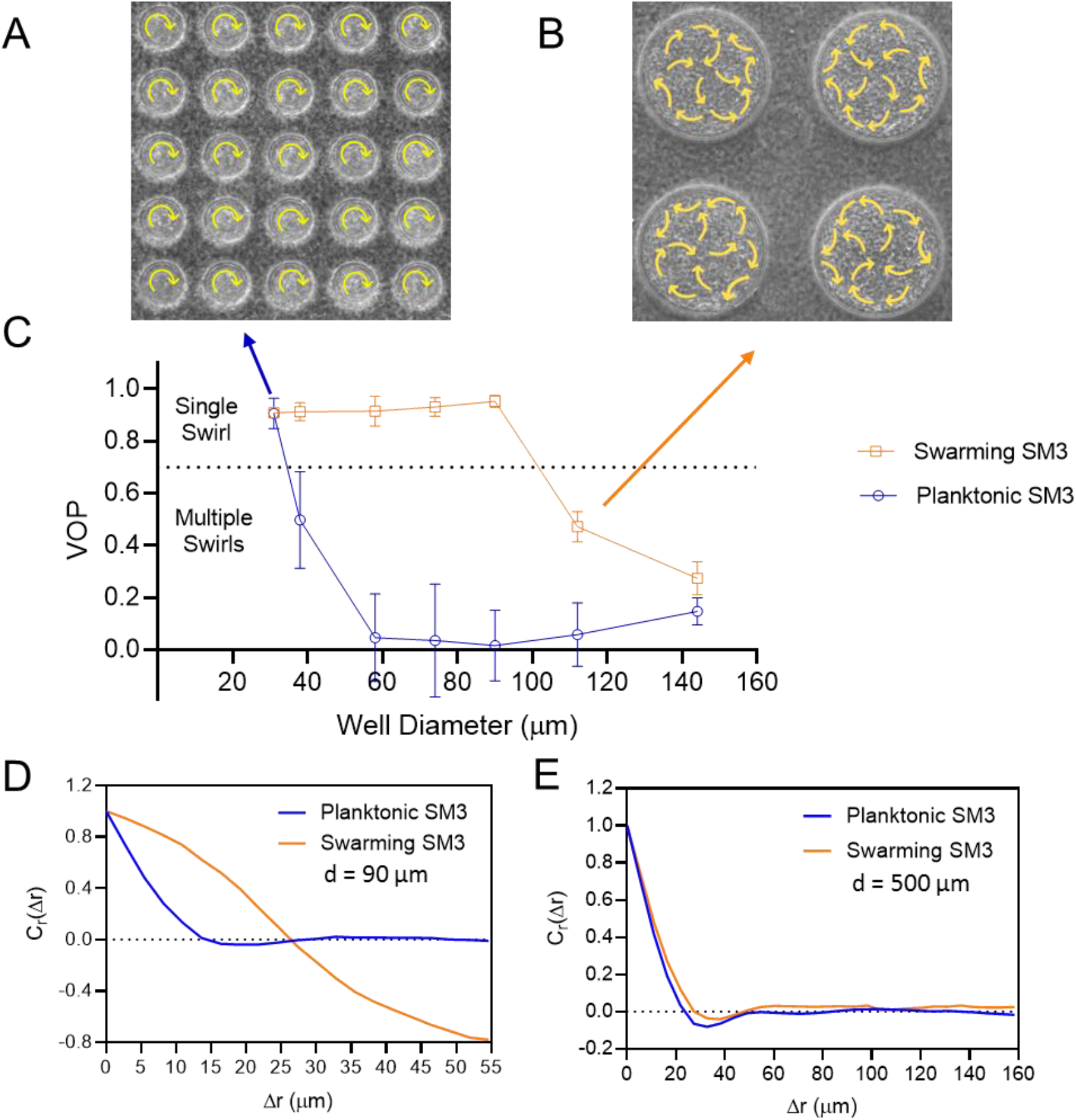
The effect of well diameter on confined *Enterobacter* Sp. SM3 motility patterns. (A-B) Motion pattern of concentrated planktonic SM3 confined in 31 μm (A) and swarming SM3 confined in 112 μm (B) diameter microwells. (C) VOP of swarming and concentrated planktonic SM3 as a function of well diameter. The error bars represent the standard deviations (± SD) for each data point, and the sample size is n = 5. (D-E) Spatial autocorrelations of the bacterial velocity field in the well diameter of 90 μm (D) and 500 μm (E). Unless otherwise noted, the depth of the wells is 22 μm.

We also tested other bacteria such as *Enterobacter* sp. SM1(Chen et al., 2020), *Serratia marcescens* (including one lab strain Db10 and another strain, H3, isolated from a human patient)(Chen et al., 2020), *Citrobacter koseri* (H6)(Chen et al., 2020), and *Bacillus subtilis* 3610(Chen et al., 2020). All the tested strains, with the exception of *B. subtilis,* showed similar motion pattern divergence between confined planktonic cells and swarming cells like SM3 (Fig. S1A, see discussion).

### The large single swirl behavior is indicative of cohesive cell-cell interaction

We performed several experiments to explore parameters defining the divergence of motion patterns in confinement. First, we rule out cell density difference as the reason for the difference in the confined motion patterns by concentrating planktonic cells to a comparable density of a naturally expanding swarm on agar (see Methods) before mounting the PDMS chip. Second, we noticed that SM3 tends to get elongated when they swarm(Chen et al., 2020). We hypothesize that elongated bacteria may enhance the local alignment of the rod-shaped cells and increase the vortices’ size in mesoscale turbulence(Doostmohammadi, Adamer, Thampi, & Yeomans, 2016). Thus, we treated SM3 planktonic cells with cephalexin (CEP) which has been shown to elongate *E. coli* (Hamby, Vig, Safonova, & Wolgemuth, 2018). This treatment indeed caused the cell length of SM3 to reach that of swarming cells on average (Fig. 3A). However, we found no significant change following the centrifugation and CEP treatment of the planktonic SM3 (Fig. 3B). Although CEP treated planktonic SM3 has similar cell length, cell density, and cell speed as swarming SM3, we could not restore the single swirl pattern in 74 μm confinement wells (Fig. 3C). Third, noticing a surfactant rim on the swarming SM3 colony edge, we conjectured that surfactants secreted by swarming SM3 might help align the swarmers in confinement. As a prototypical surface wetting agent, surfactin was added in several concentrations to planktonic SM3 to test whether it could promote a single-swirl motion pattern. However, it did not establish a stable single-swirl pattern. Finally, we found that adding lyophilized swarming supernatant to swimming SM3 did not increase the VOP either (Fig. 3C).

**Figure 3.**
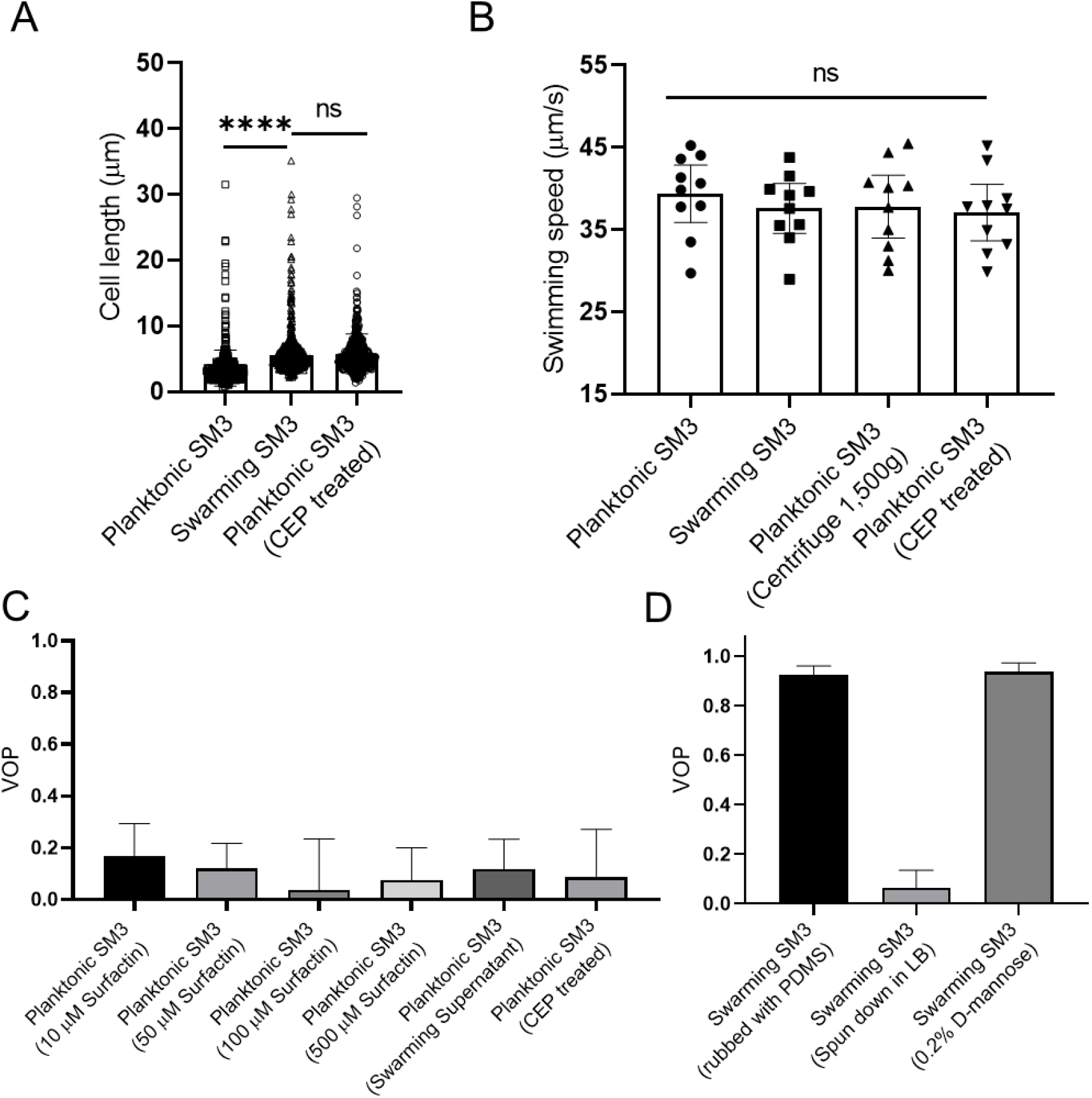
Factors that possibly influence the bacterial motion pattern in the well. (A) Bacterial cell length of planktonic, swarming, and cephalexin (CEP) treated planktonic SM3, n = 500 for each group. Data are represented as median and interquartile range. **** indicates *P* < 0.0001. ns indicates not significant (Kruskal-Wallis test). (B) Bacterial cell speed of swimming, swarming, centrifuged, and CEP treated swimming SM3, n = 10 for each group. ns, not significant, one-way ANOVA followed by Tukey’s post hoc test. (C) VOP of swimming SM3 under 74 μm diameter confinement with different treatments, n = 5 for each group. (D) VOP of swarming SM3 under 74 μm diameter confinement with different treatments, n = 5 for each group. B-D, Data are represented as mean and standard deviation (±SD).

Unable to make the concentrated planktonic SM3 form a single swirl in the 74 μm well, we tackled the problem from another angle, by altering the conditions of swarming SM3 in order to break the single swirls. Initially, we tried to physically “disrupt” the swarming colony by rubbing the swarming colony gently with a piece of PDMS offcut. This operation did not break the single swirl pattern in the wells (Fig. 3D). Then, 0.2% D-mannose was added to the swarming colony to de-cluster bacteria bundles due to cells’ sticking to each other(Hamby et al., 2018). However, this treatment could not alter the single swirl pattern, either (Fig. 3D). Finally, we diluted the swarming cells in Lysogenic Broth (LB) by 20-fold. After re-concentrating the cells by centrifugation and removing extra LB to recover the initial cell density, these “swarming” SM3 cells were pipetted back on the agar plate. After this treatment, the previous single swirl turned to multiple swirls under the confinement (Fig. 3D), suggesting that these cells now behave much like planktonic cells. We conclude that the single swirl pattern depends on cohesive cell-cell interaction mediated by biochemical factor/s removable by matrix dilution.

### Diluted swarming SM3 show unique dynamic clustering patterns

We suspected that specific interactions between the neighboring swarming cells were weakened or diminished upon dilution with the LB medium. A fifty (50) μL water droplet was applied to the swarming and the concentrated planktonic SM3 colony edges to investigate the potential intercellular alignment at a microscopic scale within the bacterial colony. In the diluted swarming colony, groups of cells formed bacterial rafts, a characteristic feature previously associated with gliding motility(Be'er & Ariel, 2019; Kearns, 2010a). Those cells within a polar cluster are moving in the same direction in a cohesive pack at the same speed (Movie S3). In contrast, upon dilution of the concentrated planktonic SM3, the cells disperse uniformly, and their moving directions appear random (Movie S4). Swarming SM3 cells tend to move together near the agar surface, while planktonic SM3 cells swim freely in the bulk fluid (Fig. 4A-B). We used the MATLAB PIV toolkit to track the moving bacteria in the image sequences of diluted swarming and planktonic SM3 for comparison. We found that swarming SM3 formed clusters with more than 20 cells on average, while we did not see such clusters of planktonic SM3 cells (Fig. 4C-D). The lingering clusters of cells in the swarming phase upon dilution point to a more substantial cell-cell cohesive interaction than between planktonic cells.

**Figure 4.**
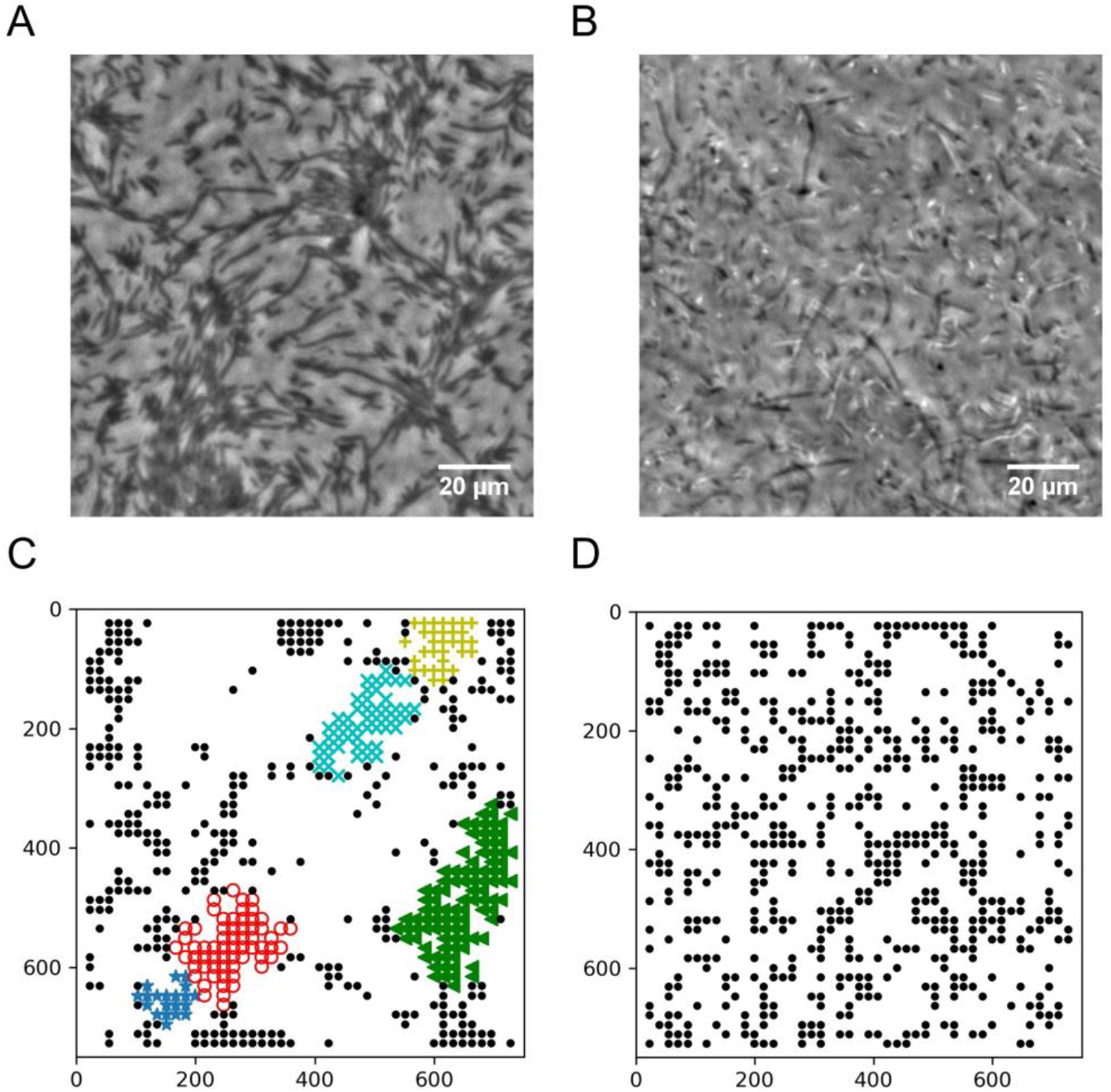
Spatial distribution of swarming and swimming SM3 cells. (A-B) Snapshots showing diluted swarming SM3 (A) and planktonic SM3 (B) on a soft agar surface, respectively. (C-D) DBSCAN clustering analysis of diluted swarming SM3 (C) and planktonic SM3 (D). Black dots represent moving bacterial cells and colored markers show cells in clusters, as determined by the program. The axis represents the dimension of the image in pixels.

### Numerical simulation reveals cell-cell interaction to be the key player

To further verify that rafting in swarming is a crucially relevant factor to the motion pattern discrepancy, we performed computer simulations using a zonal model for pair-wise interactions. The interactions among the moving particles (short-range repulsion, velocity alignment, and anti-alignment) are considered, all as functions of the particle-particle distance(Grossmann, Romanczuk, Bar, & Schimansky-Geier, 2014, 2015). The particles’ speed is fixed for simplicity, but the initial particle positions and initial moving directions are randomized. In the simulations, we interpret the rafting as due to a lower repulsion force and more substantial alignment among the swarmers (described in Methods and Supporting Text). We simulated the situation of confined swarmers and planktonic cells in different sizes of circular confinement, as performed in the experiments. The simulation results mirror the experimental results. Both swarmers and planktonic cells start with a single-swirl pattern; as the circle size is increased, the planktonic cells break into a multi-swirl motion pattern earlier than the swarmers and finally both converge to the multi-swirl region (Fig. 5A, compared with Fig. 2C; also see Fig. S3 and Movie S5). We then performed the “dilution” simulation for both states, finding that swarming cells form dynamic polar clusters when the cell density is around ρ = 235. In contrast, the planktonic cells form a “gas” phase without clustering at all comparable densities (Fig. 5B, Movie S6). This result supports the experimental observation in Fig. 4A. By encoding more vital cell-cell interaction among the swarming cells, our computational model phenocopied the experimental results in both confinement and dilution experiments.

**Figure 5.**
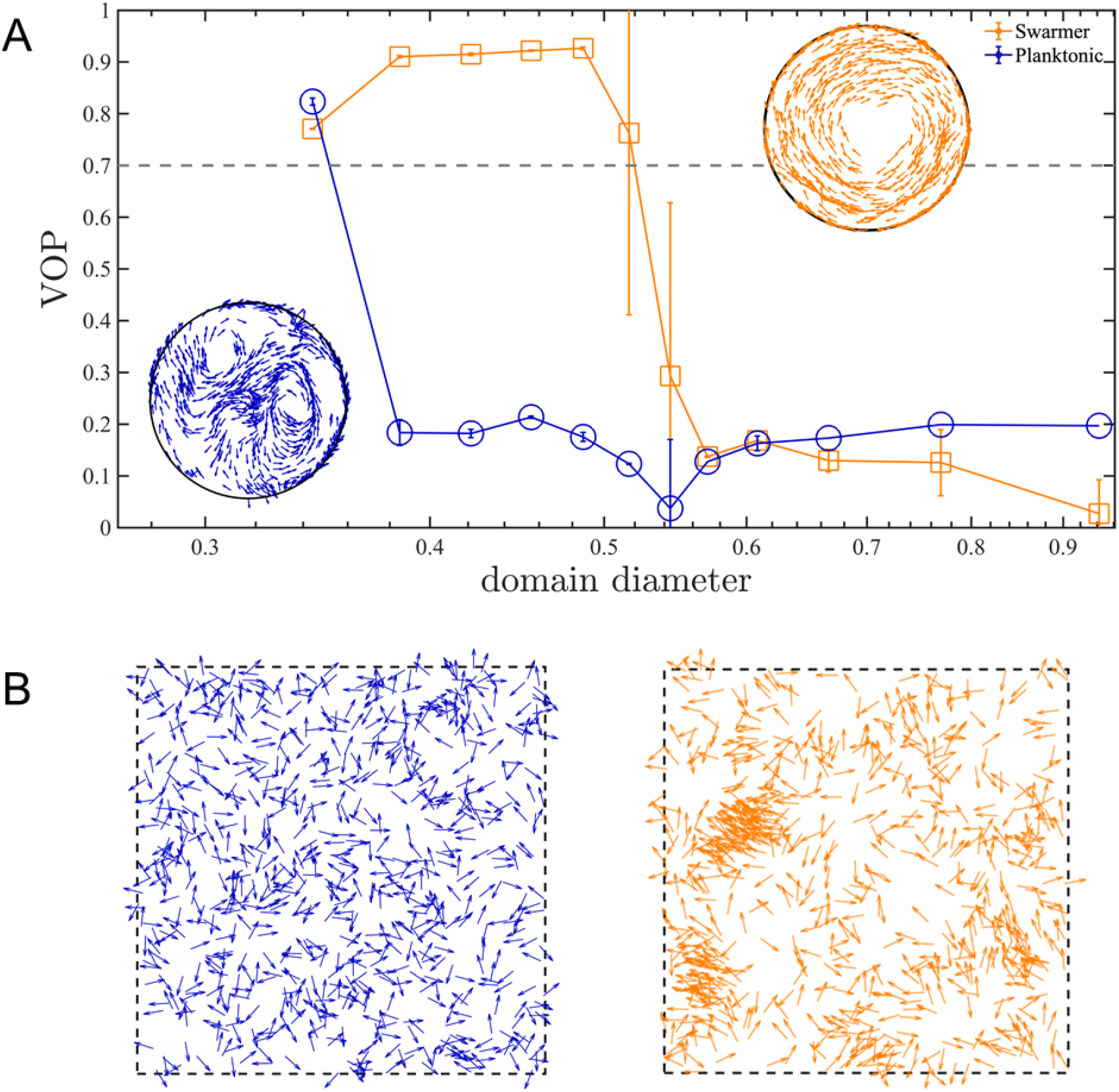
Numerical simulations of planktonic and swarming SM3 in confinement and open space. (A) VOP of swarming and concentrated planktonic SM3 as a function of well diameter. The error bars represent the standard deviations (± SD) for each data point, and the sample size is n = 5. The circles on the upper right corner and the lower left corner show representative motion patterns of swarmers and concentrated planktonic cells in the confinement size between 0.38 and 0.5. (B) Planktonic cells (left) and diluted swarming cells (right) with same cell density in a space of periodic boundary condition.

### Identifying SM3 motility type on mice mucosal surface

The difference in confined motion patterns enables us to detect bacterial swarming on surfaces other than agar, including physiological environments such as mucosal. We are unaware of any previous studies or examples regarding bacterial swarming on non-agar surfaces. There are considerable technical challenges in dealing with uneven or more complex surfaces. The mouse intestinal tissue, for instance, is more than 1 mm thick and non-transparent. Since light cannot penetrate the tissue, observing bacteria directly on the tissue surface is not feasible. Staining or fluorescence labeling may alter the bacterial swarming motility (e.g. we found that SM3 becomes non-swarming once GFP labeled). If the bacterial cells are labeled biochemically, the fluorescence signal weakens when the cells reproduce. As an alternative strategy, using PDMS chips coated with fluorescent beads and then mounted on SM3 inoculated C57BL6 mouse intestine tissue, we detected swarming motility based on the “collective” swirling motion of the beads (see Methods, Fig. 6, and Movie S7&8). This experiment on the mouse intestine tissue confirms that bacterial swarming indeed occurs on a non-agar, physiologically relevant surface.

**Figure 6.**
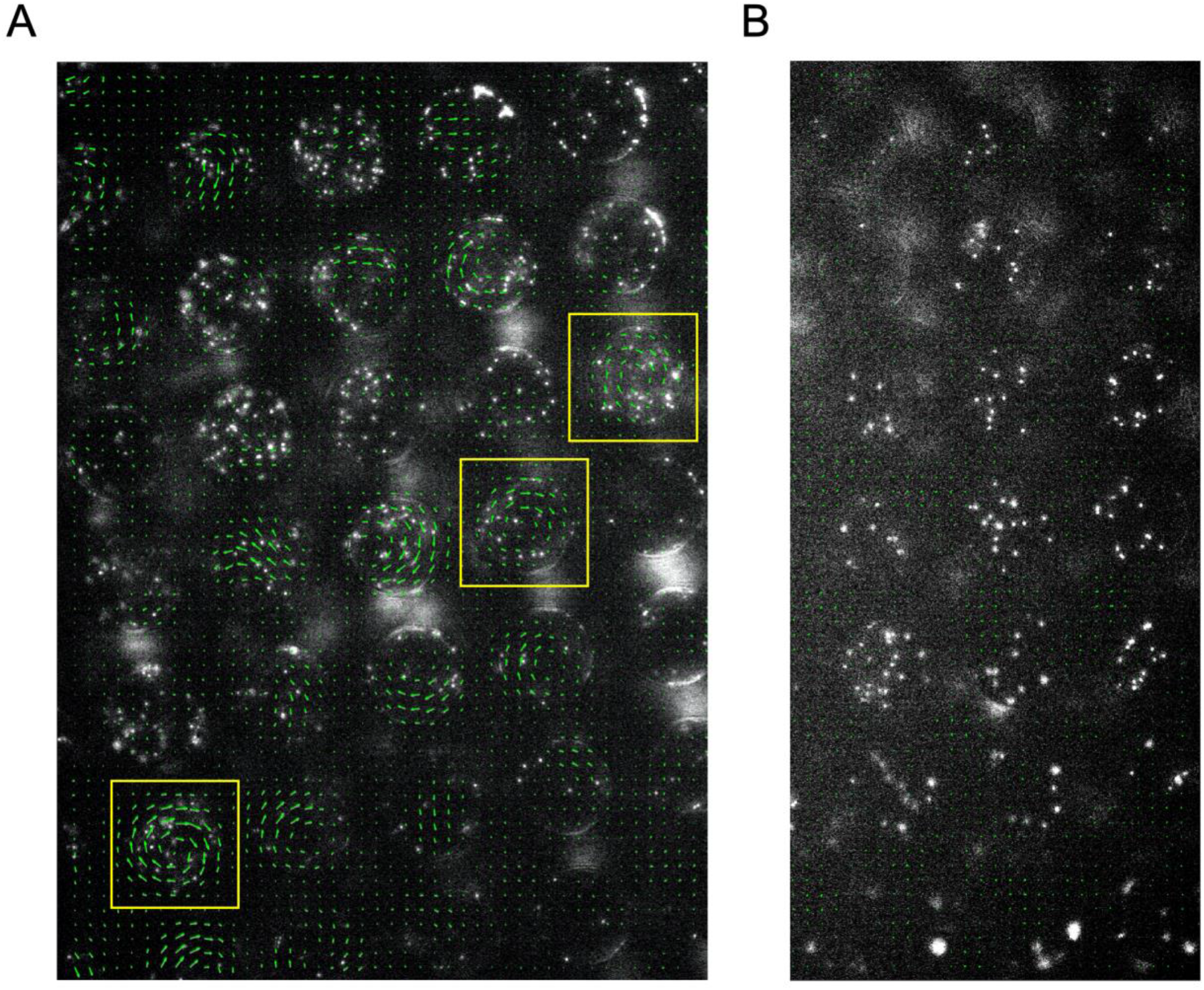
Motion of fluorescent beads in microwells mounted on infected murine tissue. PDMS chips were coated with 0.5 μm fluorescent beads and mounted on SM3 inoculated colitic (A) or non-colitic (B) mice intestine tissue surfaces. The beads motion was measured after 4.5 hr incubation. Average velocity field was calculated by tracing the beads motion using PIV toolkit. (A) On colitic tissue, wells with VOP > 0.7 were found and marked with yellow squares. We conclude that, in these wells, the single swirl motion pattern of the beads was powered by the confined swarming SM3. Since the tissue surface was not as smooth as on agar surface, the motion of the beads in some wells did not form a complete vortex, yet jets indicating partial vortices were discernable. (B) On a normal tissue lacking swarming bacteria, the average velocity of the beads in the wells due to random motion was close to zero, giving rise to uniformly small VOP values. We could infer that the confined SM3 in these wells were predominantly swimming rather than swarming.

## Discussion

We have shown the motion pattern differences between PDMS chip confined planktonic and swarming *Enterobacter* sp. SM3 in the size range of 40 μm ≤ d ≤ 90 μm. Compared with previous work, our experimental setup has the advantage of ensuring stable and sustainable patterns. First, PDMS material does not harm living bacteria cells and is permeable to oxygen(Turner, Zhang, Darnton, & Berg, 2010), thus ensuring continued oxygen exposure required for swarming(Chen et al., 2020). Second, we mounted the microchip on a soft agar containing over 97% water, which automatically fills the wells via permeability and capillary flow. Finally, the LB agar also provides the necessary nutrients to fuel the bacterial movement in the wells. Therefore, bacterial cells confined in the microwells remain motile for hours, much longer than in droplets surrounded by mineral oil(Hamby et al., 2018; Wioland et al., 2013) or in microfluidic chambers with glass surfaces(Beppu et al., 2017; Wioland, Lushi, et al., 2016), where bacteria movement typically lasted no more than 10 minutes.

Prior studies have proposed different models to explain the circularly confined motion of rodshaped swimmers(Hamby et al., 2018; Lushi et al., 2014; Tsang & Kanso, 2015). However, previous models cannot explain the motion pattern difference we observed for confined swarming and planktonic SM3. Noticing that swarming SM3 washed in LB lost the single swirl pattern, we hypothesize that other than cell length or cell speed, the strong cell-cell interaction may be a key factor responsible for the persistence of single swirls in the wells. The mechanism of the rafting phenomenon of swarming cells has not been fully deciphered yet(Kearns, 2010a). It might be due to cohesive interaction among neighboring cells and hydrodynamic effects among 2D-confined peritrichous flagellated bacteria(Li, Zhai, Sanchez, Kearns, & Wu, 2017). The cell-cell interaction may further result from biochemical change of cell envelope during swarming (e.g., more long sidechain lipopolysaccharides) or secretions(Armitage, Smith, & Rowbury, 1979). Once these surrounding matrix or polymers are washed away by ~ 100-fold dilution, the cohesive interactions are diminished, resulting in a loss of dynamic clusters in the dilution experiment, and multi-swirl motion pattern under confinement. We confirm that lower repulsion and higher alignment are the key factors that differentiate swarmers and planktonic cells by reproducing the experimental results via numerical simulation. Future work is called upon to explore the swarmer rafting phenomenon further and investigate the molecular basis for cell-cell cohesive interaction among the swarming cells.

A spectrum of swarming bacteria manifests the same characteristic as SM3 (Fig. S1A). The bacteria tested, including SM1, H6, H3, and Db10, all behave like SM3. They all showed clustering or cohesive cell-cell interaction when the swarming colony was diluted and uniformly dispersed when the concentrated planktonic cells were diluted. One notable exception is *Bacillus subtilis.* Swarming and concentrated planktonic *Bacillus subtilis* 3610 show the same motion pattern across different confinement sizes. For well diameter d ≤ 90 μm, both swarming and swimming *B. subtilis* form single swirls while for well diameter d ? 112 μm, they both break into mesoscale vortices. *B. subtilis* is a Gram-positive bacterium, different from SM3, SM1, H6, H3, and Db10. We speculate that swarming *B. subtilis* does not have as strong a cell-cell interaction as SM3 and its gram negative cohort we tested. The interaction is not so different between the swarming and planktonic *B. subtilis* 3610 cells since we found the diluted swarming cells to disperse uniformly, and with no clustering behavior, much like diluted planktonic cells. The swarming colony thickness for *B. subtilis* may also play a role in defining the differences between this bacterium and the other strains. It is known that swarming *B. subtilis* produces abundant surfactant, resulting in a wide-spread, monolayer, non-compact colony(Be'er & Ariel, 2019; Jeckel et al., 2019). In contrast, swarming SM3 and the other tested bacteria are multilayer colonies that can be as thick as 20 - 40 μm. The thickness of SM3 swarm and that of its gram-negative cohort on agar may extend the strong cell-cell alignment through the entire depth of PDMS wells, which is lacking among planktonic cells of comparable concentration (Fig. S1B).

Our experiments on SM3 confirm the prediction made by Beppu *et al.* that single vortex occurs when the confinement diameter d is smaller than a critical length *l*\*(Beppu et al., 2017). Here, the critical length for swarming SM3 is ~ 49 μm, whereas, for concentrated planktonic SM3, it is ~ 17 μm. Interestingly, the same bacterial strain in different motility states has two distinct critical lengths. Thus, we were able to use this property to identify the motility types on mouse mucosal surfaces. The beads’ motion is not a perfect swirl in every well on the colitic tissue because the mucosal surface is not as smooth as the agar surface. There are sags and crests on the inflamed mucosal surface due to the disrupted mucin layer(Chen et al., 2020). We conjectured that this unevenness hindered the swirl formation to a certain extent. Indeed, intact swirl patterns were spotted only on limited locations where the mucosal surface was relatively flat. Nevertheless, capturing only a few wells where beads showed single swirl motion was sufficient to show that swarming occurred on a mucosal surface.

Evidence of genetic and epigenetic regulation(Daniels, Vanderleyden, & Michiels, 2004; Morgenstein, Szostek, & Rather, 2010; Tremblay & Deziel, 2010; Wang, Frye, McClelland, & Harshey, 2004), and cell morphology changes (e.g., cell elongation and hyper-flagellation), indicates that swarming is a different phenotype from swimming. Lacking comparison under the same experimental conditions, one might suspect that bacterial swarming might be a dense group of cells swimming on a surface(Kearns, 2010a). Here, through geometry confinement, we show that *Enterobacter* sp. SM3 swarming manifests different biophysical characteristics from swimming. This study’s key experimental method differentiates swarming motility from swimming motility at mesoscopic or even macroscopic scales, providing a visual assay to detect swarming behavior on either an agar or tissue surface. This study’s findings provide the rationale for developing applications such as isolating bacterial swarmers from a polymicrobial environment and developing diagnostics for the presence of *in vivo* swarming (e.g., detecting urinary or fecal swarming bacteria in catheter infections or intestinal inflammation, respectively)(Arikawa & Nishikawa, 2010; Lane et al., 2007). Additionally, the sensitivity to confinement size indicates that a quantitative ranking system for different swarmers could be established based on the characteristic well size that stabilizes the confined motion pattern into a single swirl. Such a ranking system will be significant for future investigations on the implications of swarming bacteria in host physiology and pathophysiology.

## Methods

### PDMS confinement sheet fabrication

Polydimethylsiloxane (PDMS) microwell confinement sheets with different combinations of well sizes and depths were fabricated using a soft photolithography technique. Patterns of the confinement were first designed using the software “L-Edit” and then uploaded into a maskless aligner (MLA 150, Heidelberg). On a 3.5-inch silicon wafer (University Wafer Inc.), photoresist gel SQ25 (KemLab, Inc.) was spin-coated at 2,000 rpm (spin speed varies according to the desired coating thickness). After baking, UV exposure, and chemical development, the microwells’ designed pattern was shown on the wafer (molding). Then, PDMS (Dow Corning Sylgard 184) base elastomer was mixed with the curing agent at the ratio of 10:1 in weight. The mixture was cast onto the patterned silicon wafer. Two grams of the mixture ended up with a PDMS sheet about 0.5 mm thick. The PDMS solidified at room temperature within 48 hours and it was cut into pieces and peeled off from the silicon wafer before use (demolding).

### Bacterial growth and confinement (Fig. 7A)

**Figure 7.**
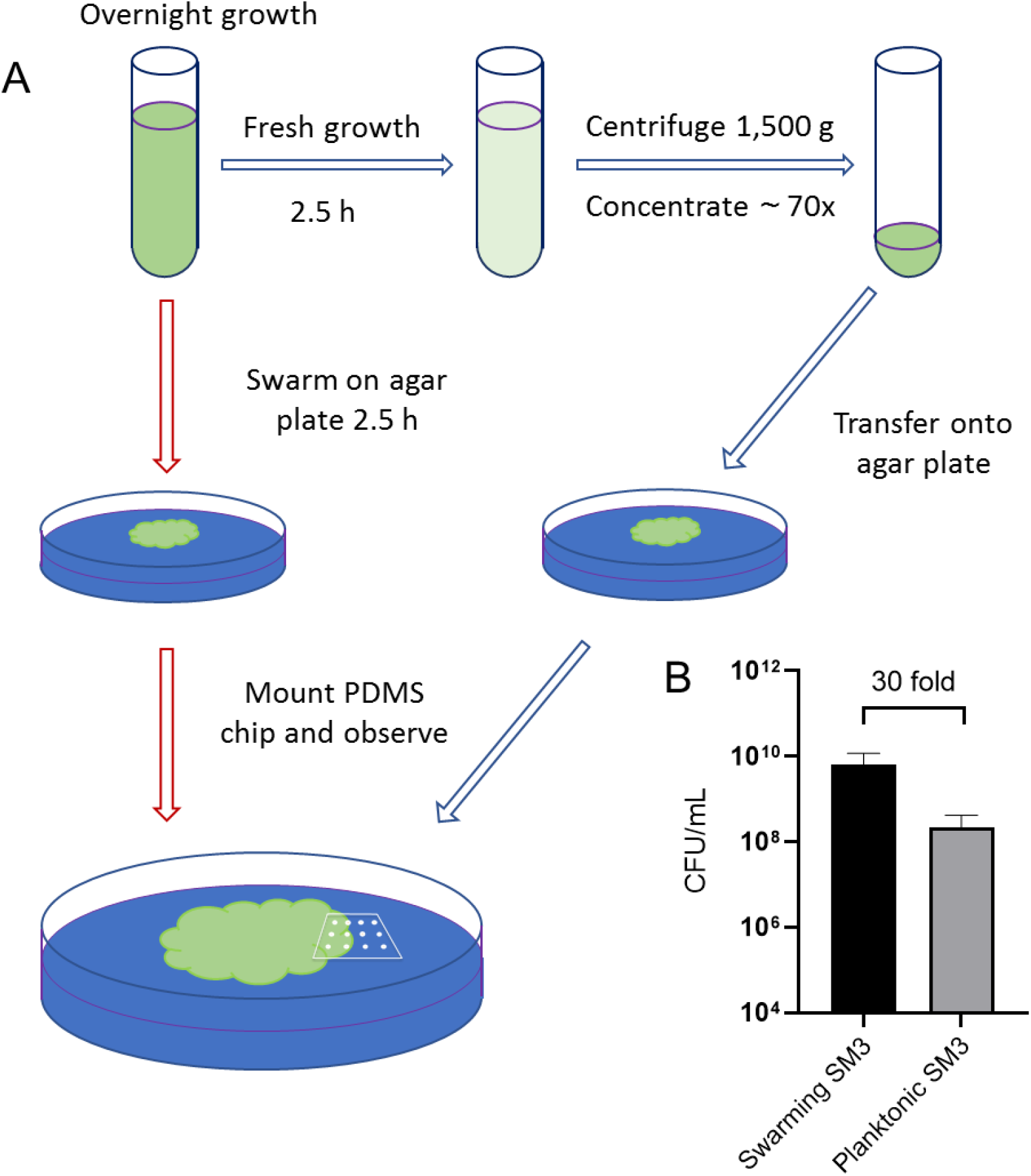
Illustration of experimental procedure. (A) Schematic of sample preparation procedure. Red arrows represent the assay procedure for swarming bacteria. Blue arrows represent the assay procedure for swimming planktonic bacteria. (B) Cell density measured by colony forming unit (CFU/mL) of swarming SM3 and swimming SM3. Swarming SM3 cell density is measured after SM3 swarming on an agar surface for 2.5 h while swimming SM3 cell density is measured for overnight SM3 culture being regrown in fresh Lysogeny Broth (LB) for 2.5 h. Since cell density of swarming SM3 was higher than that of planktonic SM3, the latter was concentrated before being applied on the agar plate to acquire comparable cell density.

*Enterobacter* sp. SM3 is a novel swarming bacterial strain isolated from inflammatory mice(Chen et al., 2020). SM3 was transferred from −80°C glycerol stock to fresh LB (Lysogeny Broth: water solution with 10 g/L tryptone, 5 g/L yeast, and 5 g/L NaCl) and shaken overnight (~ 16 h) in a 37°C incubator at 200 rpm. For swarming under confinement assay (Fig. 7A, red arrows), two (2) μL overnight bacterial culture was inoculated on the center of an LB agar plate (10 g/L tryptone, 5 g/L yeast, 5 g/L NaCl, and 5 g/L Agar; volume = 20 mL/plate) and kept in a 37°C incubator. After 2.5 h of swarming, a PDMS chip (~ 1 cm^2^) was mounted upon the edge of the swarming colony and the Petri dish was transferred onto the microscope stage for observation. For swimming under confinement assay (Fig. 7A, blue arrows), overnight bacterial culture was resuspended in fresh LB (1:100 in volume) and shaken in the 37°C incubator at 200 rpm for 2.5 h. The freshly grown culture was centrifuged at 1,500 g for 10 min and ~ 98.6% of the supernatant was removed so that the resultant cell density is about 70 times the freshly grown culture. Ten (10) μL concentrated bacteria culture was inoculated on the LB agar plate, and the PDMS chip was mounted immediately. The plate was then transferred onto the microscope stage for observation. For other bacteria strains, including *Bacillus Subtilis* 3610, the procedure was the same as that of SM3. There are thousands of wells on one PDMS chip, and when mounted on a bacteria spot or colony edge, hundreds of them are occupied by bacteria. The PDMS chip was first brought to contact with the bacteria and then gently mounted onto the agar. By doing so, there was a cell density gradient across an array of wells, with the wells closer to the bacteria spot or colony center having relatively higher cell density. We focused on the area where the confined bacteria showed collective motion, i.e. the cell density was not too high to oversaturate the well, or too low so that each cell was moving independently.

### Bacterial cell density measurement (Fig. 7B)

Two and half hour (2.5 h) freshly grown SM3 was subjected to different factors of dilution in LB, such as 10^2^, 10^3^, until 10^8^. Fifty (50) μL of each diluted culture was inoculated and spread on 1.5 % LB agar plate (10 g/L tryptone, 5 g/L yeast, 5 g/L NaCl, and 15 g/L Agar; volume = 20 mL/plate) and was incubated at 37°C for 16 h. Bacterial colonies appeared on the agar plates and the number of colonies was counted for the dilution that resulted in the colony’s number on the order of 100. The colony forming unit per microliter (CFU/mL) was calculated by dividing the colony number by the sampled volume. For swarming SM3, the cell density was measured similarly. On the edge of the swarming colony, a chunk of swarming SM3 (~ 1 mm wide) was picked by an eight (8) mm-wide square spatulate containing a small agar bottom to ensure all the cells in that region were sampled. The 1 mm x 8 mm chunk of swarming SM3 was then mixed into 1 mL LB for CFU determination. The colony thickness was assumed to be uniform across the sample. It was measured by microscopy focusing on the top of the colony and the top of the agar surface (i.e., at the bottom of the colony), keeping track of the fine adjustment knob readings. Particles of baby powder (~ several micrometers in diameter) were spread on the swarm colony surface and the agar to aid in the microscope focus. The thickness of the swarming colony was calculated based on the calibration of the knob turning tick readings.

Then the cell density was estimated by CFU/mL. CFU was calculated for both swarming and swimming SM3 to ensure the cell densities of these two cases were comparable inside the wells. We consider colony-forming unit counting a better way to control the live cell number than merely using the volume fraction because 1), Dead cells that count in the volume fraction will not contribute to the motion in the well, but they will be excluded in CFU calculation; 2), It is challenging to measure the volume of dense bacterial suspension using pipetting method due to high viscosity.

### Bacterial cell length and motility

For swimming SM3, 2.5 h freshly grown culture was diluted 100 times in LB and 50 μL of which was transferred on a glass slide and covered with a coverslip. The sample slide was placed under the microscope (Olympus CKX41, 20X), and image sequences were captured. Cell lengths were measured using ImageJ (v1.59e) freehand label tool. Cell speed was calculated as the traveling trajectory length divided by the traveling duration (~ 1s). For swarming SM3, a chunk of swarming bacteria was collected from the swarming colony edge and mixed with 1 mL LB. A glass slide and a cover slip sandwiched a droplet of 50 μL mixed culture, and the rest of the procedure was the same as that for the swimming SM3.

### Swimming SM3 with different treatments

**i), Cephalexin treatment.** Overnight SM3 culture was diluted 100 times in fresh LB and incubated in a 37°C shaker at 200 rpm for 1.5 h. Cephalexin (CEP) (C4895; Sigma-Aldrich) was added to the culture so that the CEP’s resultant concentration was 60 μg/mL. The culture was kept in the shaker for another two (2) h before use. **ii), Surfactin additions.** After 2.5 h regrown culture was centrifuged, more supernatant was removed than usual, and surfactin (S3523; Sigma-Aldrich) was added so that the resulting concentrations of surfactin were 10, 50, 100, 500 μM. At the same time, the cell density remained comparable to that of swarming SM3. **iii), Addition of swarming supernatant.** Before swarming SM3 covered the plate, the colony was scratched carefully using a PDMS (~ 0.5 cm^2^) and transferred into 1 mL deionized water. The mixture was sucked into a syringe and filtered with a 0.2 μm filter. The solution was then lyophilized to powder and then dissolved into the concentrated planktonic SM3 of roughly the same volume as the collected swarm fluid. Thus, the concentration of the swarming supernatant was kept the same to subject the concentrated planktonic SM3 to.

### Swarming SM3 with different treatments

**i), Soft scratching with PDMS.** After SM3 swarmed on the agar plate for 2.5 h, a piece of PDMS (~ 0.5 cm^2^) was used to softly scratch the edge of the swarming colony so that the swarming cells were disturbed. A PDMS confinement chip was then mounted on the disturbed region for observation. **ii)**, **Spun down in LB.** After swarming for 2.5 h, SM3 cells were collected from the colony’s edge using the blotting method(N. Darnton, Turner, Breuer, & Berg, 2004). The cells were blotted by a piece of spare PDMS and transferred to 1 mL LB. The swarming cells were centrifuged at 1,500g, and LB was removed to restore the initially high cell density. Ten (10) μL of the swarming cells thus treated were inoculated on a new swarm agar and a PDMS confinement chip was mounted for observation. **iii), D-mannose.** A droplet of 50 μL 0.2% (w/v) D-mannose (Cas No. 3458-28-4; RPI) was pipetted on a swarming SM3 colony edge. After 1-2 minutes, when the cell density became uniform again, a piece of PDMS confinement chip was applied to the D-mannose treated region for observation under the microscope.

### VOP measurement and spatial autocorrelation function

Image sequences of swarming or swimming SM3 under confinement were taken by a microscope camera (ThorLabs, Kiralux CS505MU) and then processed using a particle image velocimetry (PIV) package in MATLAB. The velocity field was marked for the confined bacteria and the VOP was calculated using the equation in Fig. 1E. Using the velocity field information, we calculated the spatial autocorrelation function through the equation 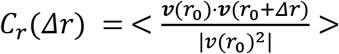, where r_0_ is the local position vector and Δr is the displacement vector(Patteson, Gopinath, & Arratia, 2018). A Python script was written to calculate all the C_r_ values in the region of interest (ROI) with a label of Δr values. These Cr values were then plotted as a function of Δr.

### Clustering analysis

On the swarming SM3 colony edge or concentrated swimming SM3 inoculation, a droplet of 50 μL deionized water was added via a pipette. Once the fluid flow stabilizes, image sequences were captured at the diluted swarming or planktonic SM3 sample locations. In a region of 130 μm x 130 μm, the velocity field was calculated using the PIV toolkit, and the vectors with magnitude below four (4) μm/s were removed. The purpose of the vector validation was to exclude non-motile bacteria. Once the moving cells were identified, a Python script was implemented to perform the clustering analysis using the function of DBSCAN(Ester, 1996) where the parameter ε was set to 50, which specifies how close points should be to each other to be considered a part of a cluster, and the minimum number of points to form a cluster was set to 20.

### Numerical Simulations

The numerical simulation consists of a 2D system of *N* particles. The position **r** of each particle is modeled via the following overdamped Langevin equation:

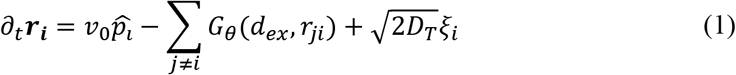

It is assumed that particles are cruising at a constant speed of *v*_0_ in the direction of 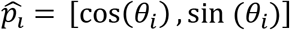. The second term includes the exclusion forcing term from all neighboring particles residing at a distance *j* closer than the exclusion range *d_ex_*. The last term is the thermal fluctuation term with the translational diffusivity *D_T_* and a zero-mean and delta-correlated noise term ξ. The direction of motion *θ_i_* of each particle is updated by the interaction terms *F_θ_*, which includes alignment, anti-alignment and repulsion effects with all neighboring particles and the rotational diffusion term with diffusivity of *D*_r_ and noise term *ζ*:

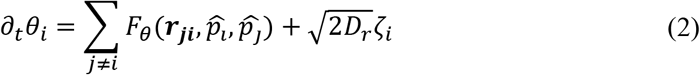

The details of the binary interaction terms *G_θ_* and *F_θ_* are provided in the Supporting Information. The simulation starts with random initial position and orientations, followed by numerical integration of equations (1) and (2) using a first-order Euler method. The integration time step Δt is chosen small enough to ensure numerical stability and independence of long-term dynamics from the time step increment. The interaction of particles with a circular bounded domain is modeled through a reflective boundary condition. The particles are reflected off the boundary with an angle equal to their incident angle. In all diluted cases, reflecting solid boundary is replaced with a periodic boundary condition to ensure that boundary scattering does not affect the dynamics in bulk.

### Detecting bacterial motility on mouse intestinal mucosal tissue using PDMS chips

Six-week-old female C57BL/6 mice (Jackson Laboratories, Bar Harbor, ME; #000664) were administered 3%(w/v) DSS (Dextran Sulfate Sodium) (MPI; # 160110) in animal facility drinking water daily to induce acute colitis(Chen et al., 2020). After 9-12 days, when the mice’s weight loss reached 20%, mice were euthanized using isoflurane anesthesia and large intestines were harvested. For controls, conventional six-week-old female C57BL/6 mice exposed to drinking water with DSS-vehicle added were also sacrificed and the intestines were collected. This study was approved by the Institute of Animal Studies at the Albert Einstein College of Medicine, Inc (IACUC # 20160706 & 00001172). Intestinal tissue was surgically exposed, cleansed with 35%(v/v) ethanol, and rinsed with PBS twice. The mucosal surface of the tissue was cultured (on gar streaks) for any residual bacteria and only used when there were no bacterial colonies on aerobic or anaerobic culture. Prior to experiments, a portion of the mucosal tissue was also harvested after ethanol cleansing for histology and to validate its histologic integrity with respect to non-cleansed DSS-exposed tissue. Tissues were spread on a 1% agar plate with the inner side facing up, and overnight SM3 bacterial culture was inoculated on one end of the tissue. The agar plate was incubated under 37°C for 4.5 hours to allow SM3 bacteria to duplicate and move on the tissue surface. PDMS chips (d = 38 μm) were coated with 0.5 μm fluorescent beads (Dragon green; Bangs Laboratory, IN) and cut into strips to fit the tissue’s size. The PDMS strip was mounted on and covered the tissue surface. Bead motion was observed under the fluorescence microscope (Olympus CKX41) with 20X objectives.

## Acknowledgment

We thank Daniel B. Kearns from Indiana University at Bloomington for providing the *Bacillus subtilis* 3610 bacteria strain, Cori Bargmann at Rockefeller University, for gifting us the bacteria strain *Serratia marcescens* Db10. We thank Hui Ma for his assistance in the cleanroom and discussion of the work. N.I.H. Grant 1R01CA222469-01 supported this work.

## Author Contribution

W.C., N.M., J.X.T., and S.M. conceived and designed the work. W.C. and N.M. performed the experiments and analyzed the data. H.K. performed the computational simulations. H.L. isolated the bacteria strains and prepared the mouse tissues. W.C., H.K., S.M., and J.X.T. wrote the paper.

## Supporting Information

**Figure S1.**
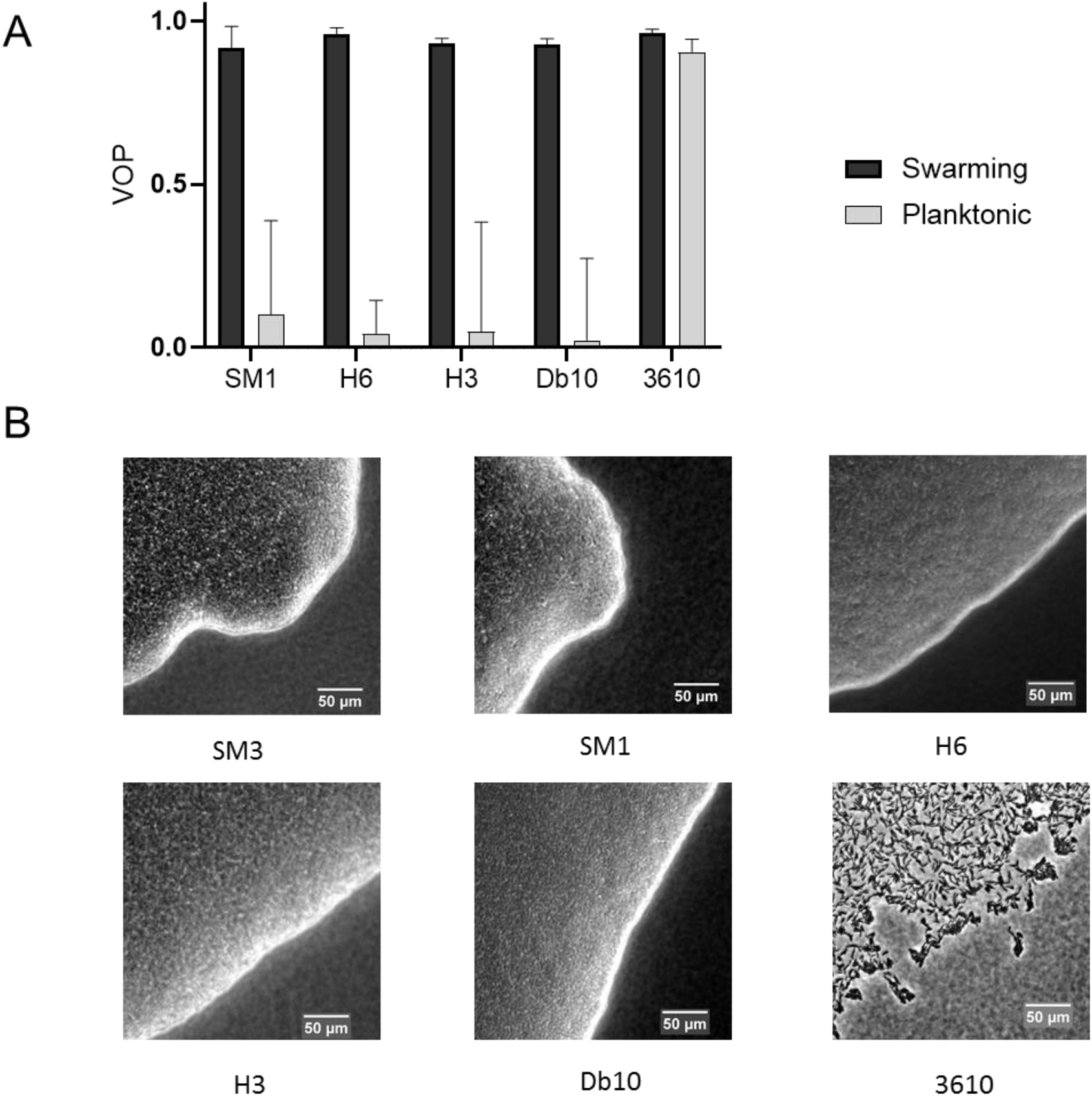
Comparison of Vortex Order Parameter (VOP) under confinement and swarm front among several bacteria species. (A) VOP of concentrated planktonic (SM1) and swarming (SM3) Enterobacter sp., *Citrobacter koseri* (H6), *Serratia marcescens* (H3), *Serratia marcescens* (Db10), and *Bacillus subtilis 3610* confined in the PDMS microwells of 58 μm in diameter and 22 μm in depth. The bars indicate averages with standard deviation (+SD) over five microwells. (B) Swarm in front of the tested bacteria. *B. subtilis* 3610 forms a monolayer, loose swarming colony while all the other bacteria strains form multilayer, compact swarming colonies.

### Supporting Text: Mathematical Modelling and Computer Simulation

#### A simplified treatment of swarming bacteria

Most particle-based models for self-propelled microswimmers incorporate detailed hydrodynamics of elongated rods in a low Reynolds number environment (Costanzo, Di Leonardo, Ruocco, & Angelani, 2012; Lushi & Peskin, 2013; Lushi, Wioland, & Goldstein, 2014; Saintillan & Shelley, 2007). However, the dynamics of bacterial swarming comprise a complex interplay between several physical and chemical interactions that go beyond hydrodynamic and steric effects. Cell interactions with the extracellular polymeric network, mechanical locking, and intertwining of flagella and formation of intercellular bundles between adjacent swimming cells (Copeland & Weibel, 2009; Kearns, 2010) are a few examples whose underlying mechanisms are not fully understood. In the absence of a comprehensive model that captures many interactions among swarming bacteria, we seek a simplified description of active particles interacting via competing interactions that capture the essential dynamics of both swarming and planktonic bacteria. Our focused aim in connection with the experimental study in this report is to discern the distinct, collective behaviors of swarming bacteria from their planktonic counterpart, in comparable concentration, and under the extent of same spatial confinement.

There are numerous approaches for incorporating the relevant physical interactions between active particles (Grossmann, Romanczuk, Bar, & Schimansky-Geier, 2014, 2015; Wensink et al., 2012; Wensink & Lowen, 2012) (readers are referred to Bär *et al.* for a recent review(Bar, Grossmann, Heidenreich, & Peruani, 2020), for example, on models for dry and wet interacting self-propelled rods). Here, we choose the binary interaction model introduced by Großmann et al.(Grossmann et al., 2014, 2015) based on the fact that hydrodynamic couplings among the swimmers can induce both alignment and anti-alignment effects(Baskaran & Marchetti, 2009). The simplified model we employ also allows us to implicitly embed the unknown interactions of cells with extracellular polymeric network and possibly, mechanical locking of flagella between adjacent cells in alignment, anti-alignment, and repulsion torque terms.

#### Numerical model and simulation

The dynamics of *N* interacting active particles have been modeled in a 2-dimensional space using the overdamped Langevin-based equations, assuming that inertia is negligible in a low Reynolds number environment. The position ***r*** and orientation *θ* of particle *i* are calculated using the following stochastic differential equations:

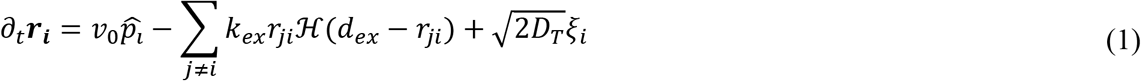

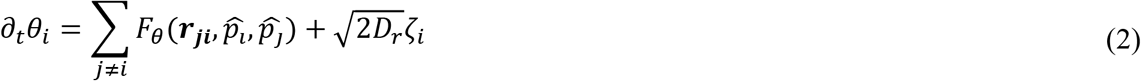

In Eq. (1), the particles’ self-propulsion speed is set to be a constant *v*_0_ along the direction 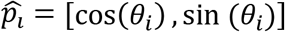. This simple assumption is based on our experimental observations, suggesting that the bacterial velocity in the suspension is largely independent of the local cell density. The second term incorporates the central exclusion force term with a spring constant *k*_ex_, which acts over the relative distance *r*_ji_ with all the neighboring particles *j*. This exclusion force term applies only when *r*_ji_ gets smaller than the exclusion range *d*_ex_ (represented as a Heaviside function 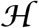). The last term in Eq. (1) is the Brownian fluctuation term with the corresponding translational diffusivity *D*_T_ and *ξ*_i_ is the white noise with zero mean and correlation *δ*(*t*).

Two terms influence the temporal change in the orientation of each particle. The first term on the right-hand side of Eq. (2) includes all the binary interaction terms. The last term on the right-hand side of Eq. (2) is the contribution from the angular Brownian fluctuation with the rotational diffusion *D*_r_ and a zero mean and delta-correlated stochastic white noise ?. In the present study, we employ the pair-wise interaction model introduced by Grossman and co-workers(Grossmann et al., 2014, 2015), which successfully reproduces various macroscopic patterns that occur in dense bacterial suspensions. The pair-wise interaction term is based on a zonal model (illustrated in Figure S2 below), capturing the alignment, anti-alignment, and repulsion effects. It is formulated in the following form(Grossmann et al., 2014, 2015):

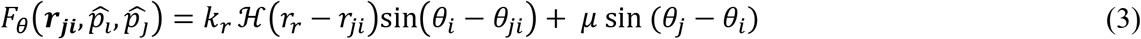

*k*_r_ is the magnitude of the constant repulsion interaction that applies over a distance of *r_r_* around the particle (Figure S2). The second term in Eq. (3) represents the alignment and anti-alignment effects, which operate over a range of *r*_a_ and *r*_aa_, respectively. The magnitude of the aligning interaction μ is distance-dependent and is defined as(Grossmann et al., 2014, 2015):

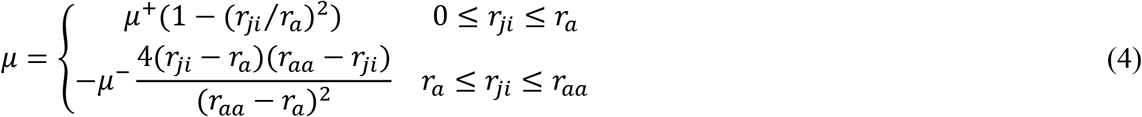

where μ^+^ and μ^-^ are the strength of alignment and anti-alignment interactions, respectively.

**Figure S2.**
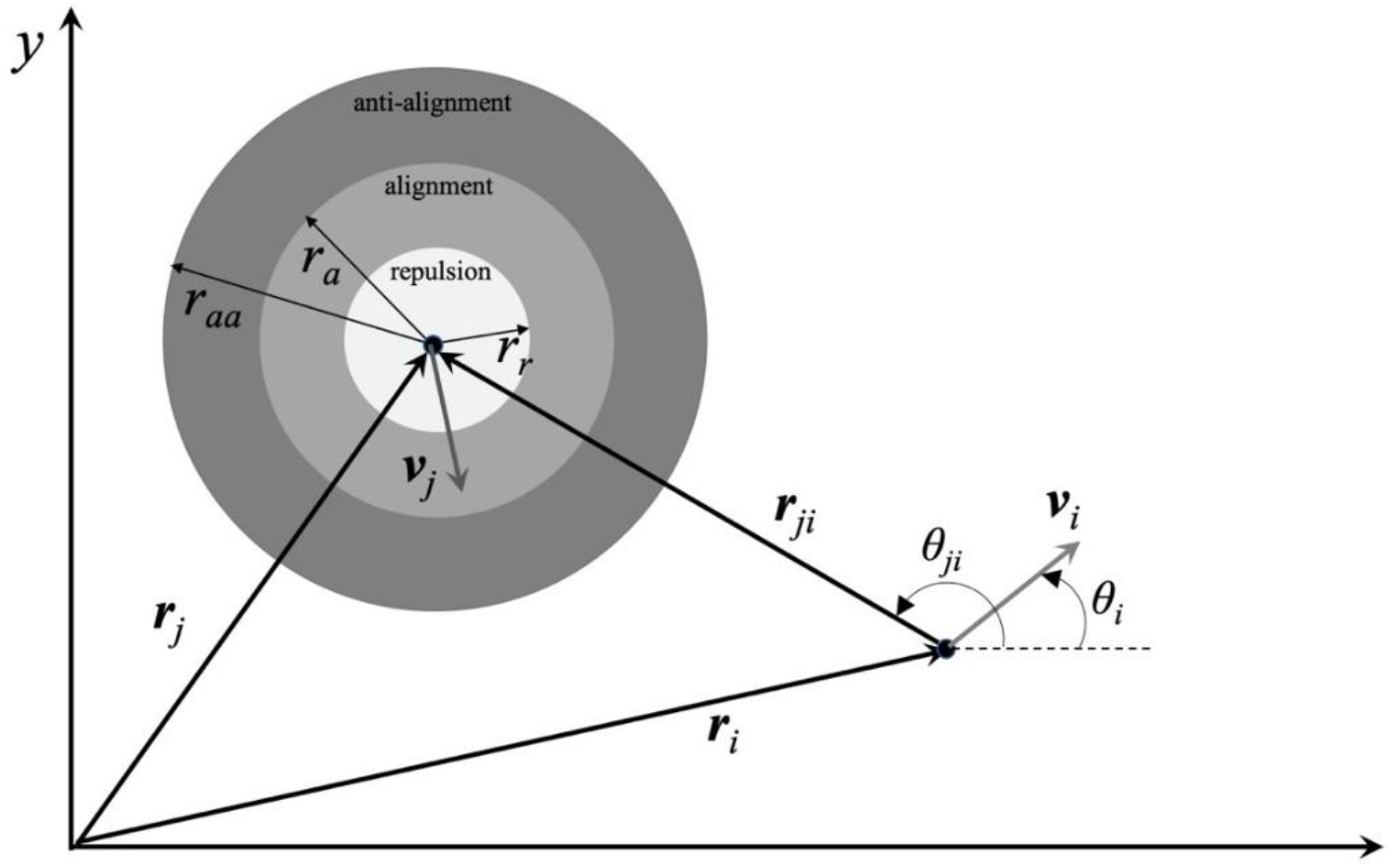
Schematic of the zonal pair-wise interaction model showing anti-alignment, alignment, and repulsion zones with the corresponding interaction radii *r*_aa_, *r*_a_, and *r*_r_.

We numerically integrate Eqs. (1) and (2) using the first-order Euler scheme. Initially, the particles are randomly distributed with random orientations. The integration time step Δ*t* is selected sufficiently small to ensure both numerical stability and also independence of long-term statistics from Δ*t*. The simulation time is set long enough to let the system reach a dynamic steady-state. The interaction of particles with the bounded circular domain is modeled via a reflective boundary condition.

#### Assessment of simulation parameters

Swarming cells secrete large amounts of surface-active compounds that modify the surface tension locally(Fauvart et al., 2012; Ke, Hsueh, Cheng, Wu, & Liu, 2015), as well as micro-viscosity of the fluid(Copeland & Weibel, 2009), which along with the formation of intercellular bundles between neighboring cells, can enhance the cohesive interaction and alignment in swarmer cells. Thus, simulation parameters must be chosen to capture different behaviors between the planktonic and swarmer cells.

Two different sets of interaction parameters have been used to differentiate the swarming and planktonic cases, and these parameters are summarized in Table S1. The values are unitless. We set the exclusion parameters *k*_ex_ and *d*_ex_ to fixed values of 0.02 and 0.035, respectively. It is also assumed that particles only experience a rotational diffusion *D*_r_ of 0.75. The simulations for both swarming and planktonic forms have been studied at two particle densities ρ = *N*/*A*_dom_, where *N* is the number of particles, and *A*_dom_ is the simulation domain area. In the high-density case, ρ = 4300, and in the dilute case, we set ρ = 235. In the dilute case, to further minimize the boundary effects, we replace the bounded domain with a periodic boundary.

**Table S1.**
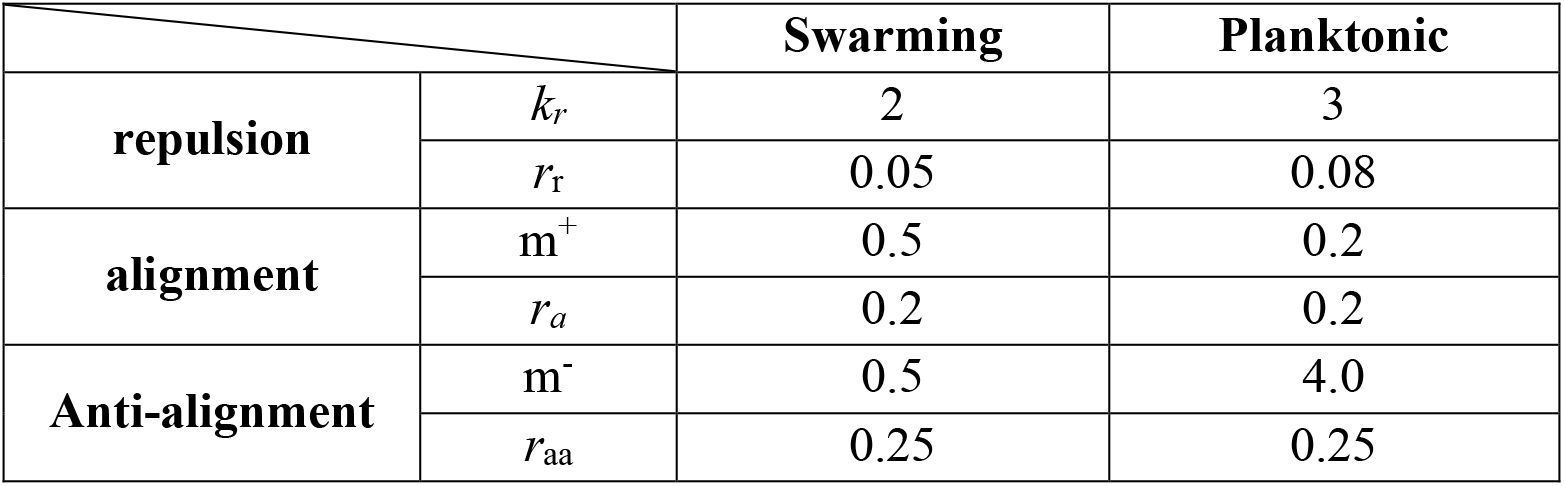
Simulation parameters used for Swarming and Planktonic cases

The simulation results at high particle density p = 4300 for some representative confinement sizes are shown in Figure S3. As Fig. S3 illustrates, the macroscopic behavior of both swarming and planktonic cells is affected by the confinement size. The corresponding change in Vortex Order Parameter (VOP) marks the transition from a single vortex to multiple swirls. Compared to the swarming case, the higher values of anti-alignment and repulsive interactions in the planktonic I case trigger an earlier onset of the transition.

**Figure S3.**
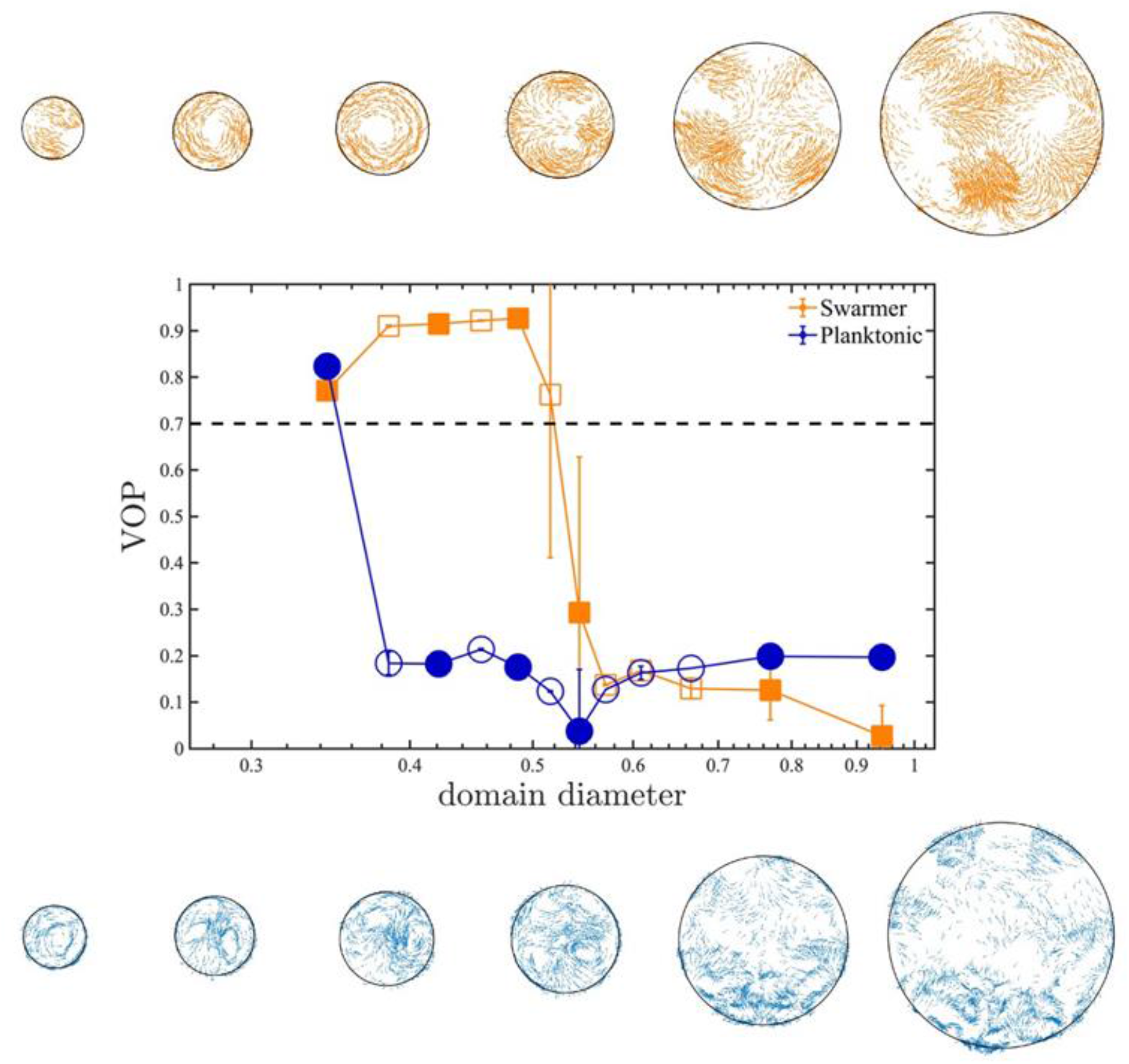
Representative patterns at different sizes of the bounded domain. Top row: Swarming; Bottom row: Planktonic. The corresponding domain sizes and VOP values are marked as filled symbols. The particle density is kept constant as the area of the simulated region increases. Simulation parameters are based on the values summarized in Table S1. ρ = 4300 in all cases.

The set of simulation parameters in Table S1 implies that (1) alignment interactions in planktonic cells are suppressed via lower alignment and higher anti-alignment magnitudes, and (2) the repulsive interaction in planktonic cells is more pronounced, in terms of higher values of the magnitude and range of repulsive torque. Despite the empirical nature of these parameter values, we found them to capture the competing interactions between planktonic and swarmer cells. The simulation results provide valuable physical insights as the patterns predicted closely resemble the experimental observation. More advanced real-time visualization of bundling dynamics in swarmer cells(Copeland & Weibel, 2009), along with biochemical characterization of the bacterial fluids, and the micro-rheology measurements within local, extracellular polymeric network(Guadayol et al., 2020) will shed light on the underlying nature of complex physical and chemical interactions. These properties rely on experimental effort beyond the scope of this report. If determined, they will facilitate the development of more comprehensive particle-based models.

### Supporting Movies

All videos play in real-time, except for Movie S7 & S8, which were taken in 20 fps but compressed to play in 30 fps.

**Movie S1: Confined swarming SM3 showing a single swirl motion pattern.** Swarming SM3 was confined in 74 μm diameter PDMS wells.

**Movie S2: Confined concentrated planktonic SM3 showing a turbulent motion pattern.** Swimming SM3 was confined in 74 μm diameter PDMS wells.

**Movie S3: Diluted swarming SM3 colony.** The swarming SM3 colony edge was diluted by adding a 50 μL water droplet. Clusters of bacteria cells formed rafts.

**Movie S4: Diluted swimming SM3 suspension.** Concentrated planktonic SM3 was diluted by adding a 50 μL water droplet. Bacterial cells were observed to swim independently without clustering.

**Movie S5: Numerical simulations of circularly confined SM3.** Swarming SM3 (left) and concentrated planktonic SM3 were simulated in the well size of 0.48. The video shows a representative confined motion pattern. Arrows indicate the moving direction of the particles.

**Movie S6: Numerical simulations of SM3 cells in open space.** Diluted swarming SM3 (left) and planktonic SM3 were simulated without confinement, but with a periodic boundary condition. In both cases, cell density is ρ = 235, and the arrows indicate the moving directions of the particles.

**Movie S7: Fluorescent beads motion on DSS induced colitic mouse intestine tissue.** The unidirectional rotation motion in 38 μm diameter wells indicates the presence of swarming SM3 on the tissue surface.

**Movie S8: Fluorescent beads motion on normal mouse intestine tissue.** The random motion in 38 μm diameter wells indicates predominantly planktonic SM3 on the normal mice tissue surface.

